# Climate Change Influences the Japanese Cedar (*Cryptomeria japonica*) Pollen Count and Sensitization Rate in South Korea

**DOI:** 10.1101/340398

**Authors:** Jung-Kook Song, Ju wan Kang, Sung-Chul Hong, Jeong Hong Kim, Dahee Park, Hye-Sook Lee, Jinho Jeong, Kyu Bum Seo, Keun Hwa Lee

**Author notes:** These authors contributed equally to this work (Jung-Kook Song, Ju wan Kang).

## Abstract

**Background:** Japanese cedar pollen (JCP) is the major outdoor allergen for spring pollinosis and seasonal allergic rhinitis (SAR) caused by JCP is the most common disease in Jeju Island, South Korea and in Japan. Prior to our research, JCP counts were strongly temperature dependent and were significantly associated with the JCP sensitization rate and JC pollinosis. This event may still be ongoing due to the effects of global climate change, such as increasing temperature.

**Methods and Finding:** For these reasons, we are studying the correlation among increasing temperatures, the JCP counts in the atmosphere and the JCP sensitization rate.

**Conclusions:** In this study, our data show that increasing temperatures in January and April might lead to earlier and longer JCP seasons and that earlier and longer JCP seasons lead to an increase in the JCP sensitization rate, which influences the prevalence of JC pollinosis.

## Introduction

Global mean air temperatures have risen at a faster rate than at any time since records began to be kept in the 1850s, and temperatures are expected to increase by another 1.8 to 5.8°C by the end of this century [1]. Climate change is expected to have enormous implications for human health, especially seasonal allergic rhinitis (SAR), such as pollinosis caused by Japanese cedar (*Cryptomeria japonica*) pollen (JCP) [2,3]. Pollinosis is among the most well-studied climate change-related diseases because pollens tend to be more active when temperatures increase and because pollens are affected by earlier and longer pollen seasons; numerous studies have shown that increasing temperatures can lead to earlier and longer pollination seasons, which result in increases in the pollen sensitization rate and in symptom duration [3].

JCP is the major outdoor allergen for spring pollinosis, and SAR caused by JCP is the most common disease in Jeju Island, South Korea, and in Japan [2, 3]. Prior to our research, JCP counts were strongly temperature dependent and were significantly associated with the JCP sensitization rate and JC pollinosis [2].

This event may still be ongoing due to the effects of global climate change, such as increasing temperature. For these reasons, we are studying the correlation among increasing temperatures, the JCP counts in the atmosphere and the JCP sensitization rate. In this study, our data show that increasing temperatures in January and April might lead to earlier and longer JCP seasons and that earlier and longer JCP seasons lead to an increase in the JCP sensitization rate, which influences the prevalence of JC pollinosis [2].

## Methods

Burkard 7-day recording volumetric spore traps (Burkard Manufacturing Co., Rickmansworth, UK) were installed in two geographic locations, 126°31’13.67” E, 33°29’29.27” N, representative of Jeju City (northern region, NR) and 126°33’18.97” E, 33° 15’11.86” N, representative of Seogwipo City (southern region, SR) (Fig. 1). The drum in the device was harvested weekly at the same time. The trapped pollen was mounted in glycerin jelly and stained. The JCP collected per 24-hour period was identified and counted by an expert at ×400 magnifications. The JCP season each year was defined as extending from the first day when JCP was detected on 2 or more consecutive days to the first day when no JCP was identified for a full day. Monthly mean temperature data for the two regions were obtained from the database of the Korea Meteorological Administration [4].

**Fig. 1.**
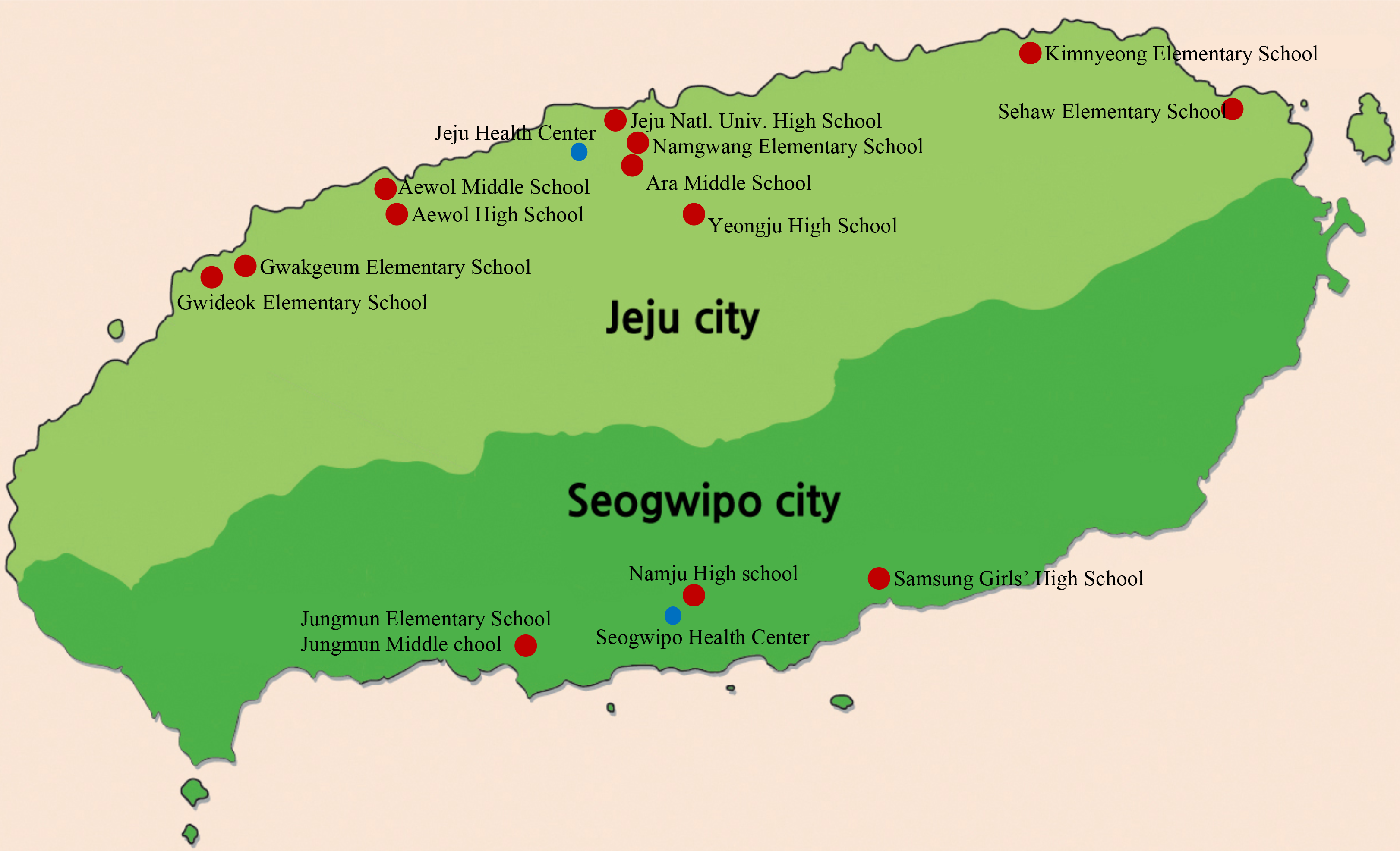
The approximate geographical locations of the selected schools (red circles) and the pollen counters (blue circle) are marked.

In Jeju Island, there are two school districts that correspond with the city districts. A balanced selection of schools and grades was made from each school district. The attendees of the schools located in Jeju City are representative of residents of Jeju City (northern region, NR), and those in Seogwipo City are representative of Seogwipo City (southern region, SR) (Fig. 1). All students in the school years 2010 and 2016 were included (Table 1). This study was approved by the institutional review board (IRB) at the Jeju National University Hospital and the skin prick test was performed after informed consent was obtained from each student’s parents. JCP (Greer Laboratories Inc., Lenoir, NC, USA) was diluted with 0.9% saline to a protein concentration of 100 μg/mL, and the same volume of 50% glycerin was added. Sensitization to antigens of JCP was defined as a mean wheal size that was the same or larger than that of the positive control (allergen/histamine ratio≥1). The data were excluded when the wheal size for the positive control (histamine, 1 mg/mL) was smaller than 2 mm or if a wheal was observed in the negative control (0.9% saline).

**Table 1.** Demographic characteristics.

|  | 2010 year | 2016 year | *p*-value |
| --- | --- | --- | --- |
| Total N (%) | 1549 (100%) | 1067 (100%) |  |
| Boys, N (%) | 763 (49.3%) | 514 (48.2%) |  |
| Girls, N (%) | 786 (50.7%) | 553 (51.8%) | 0.585 |
| By age, N (%) |  |  |  |
| 9 | 191 (12.3%) | 155 (14.5%) |  |
| 10 | 195 (12.6%) | 224 (21%) |  |
| 11 | 231 (14.9%) | 94 (8.8%) |  |
| 13 | 267 (17.2%) | 177 (16.6%) |  |
| 14 | 277 (17.9%) | 145 (13.6%) |  |
| 16 | 388 (25.0%) | 272 (25.5%) | 0.906 |
| By school, N (%) |  |  |  |
| Elementary (9 to 11) | 617 (39.8%) | 473 (44.3%) |  |
| Middle (13 to 14) | 544 (35.1%) | 322 (30.2%) |  |
| High (16) | 388 (25.0%) | 272 (25.5%) | 0.204 |
| Residential area |  |  |  |
| Jeju City (NR) | 908 (58.6%) | 543 (50.9%) |  |
| Seogwipo City (SR) | 641 (41.4%) | 524 (49.1%) | 0 |

NR, residents of Jeju City; SR, residents of Seogwipo City.

## Results

The JCP season began earlier and lasted longer each year. The JCP season was estimated from 2011 to 2017. In the NR, the JCP season lasted for 44 days in 2011, 66 days in 2012, 66 days in 2013, 67 days in 2014, 57 days in 2015, 56 days in 2016, and 71 days in 2017. In the SR, the JCP season lasted for 48 days in 2011, 73 days in 2012, 74 days in 2013, 80 days in 2014, 63 days in 2015, 69 days in 2016, and 85 days in 2017 (Table 2). Table 2 also shows that the JCP season started earlier and lasted longer in the SR than in the NR and that the level of JCP in the atmosphere in the SR was estimated to be 2-8 times higher than that in the NR (data not shown) [2].

**Table 2.** JCP season in Jeju Island.

| Area | Year | JCP detected in the atmosphere |  |  |  |
| --- | --- | --- | --- | --- | --- |
|  |  | Start | Peak | End | Duration, days |
| Jeju City (NR) | 2011 | 23-Feb | 12-Mar | 07-Apr | 44 |
|  | 2012 | 05-Feb | 01-Mar | 10-Apr | 66 |
|  | 2013 | 29-Jan | 28-Feb | 04-Apr | 66 |
|  | 2014 | 30-Jan | 29-Feb | 06-Apr | 67 |
|  | 2015 | 03-Feb | 20-Feb | 30-Mar | 57 |
|  | 2016 | 05-Feb | 26-Feb | 31-Mar | 56 |
|  | 2017 | 02-Feb | 01-Mar | 13-Apr | 71 |
| Seogwipo City (SR) | 2011 | 17-Feb | 01-Mar | 05-Apr | 48 |
|  | 2012 | 30-Jan | 03-Mar | 11-Apr | 73 |
|  | 2013 | 13-Jan | 21-Feb | 13-Apr | 74 |
|  | 2014 | 20-Jan | 02-Mar | 06-Apr | 80 |
|  | 2015 | 26-Jan | 16-Feb | 29-Mar | 63 |
|  | 2016 | 29-Jan | 08-Mar | 06-Apr | 69 |
|  | 2017 | 20-Jan | 01-Mar | 14-Apr | 85 |

During the JCP season, the mean temperatures in January and April increased each year, the increasing mean temperatures in January, March and April highly correlated with the start and end of JCP production (*p*=0.003 in January, *p*=0.897 in February, *p*=0.018 in March and *p*=0.031 in April) (Fig. 2), and the mean temperature in SR was still higher than that in NR (Table 3 and Fig. 2) [2]. The JCP sensitization rate increased year to year. Among schoolchildren in Jeju City, the JCP sensitization rate was 11.2% in 2010 and rose to 13.9% in 2016, and there was a difference in the JCP sensitization rate between the geographic regions. The JCP sensitization rate in SR was 23.6% in 2010 and 27.5% in 2016, while that in NR was 10.2% in 2010 and 13.8% in 2016. The JCP sensitization rate was higher in high school students than in students in other grades (Table 4-1, 4-2, 4-3 and 4-4). Additionally, the JCP sensitization rate was 2-8 times higher in SR than in NR [2].

**Fig. 2.**
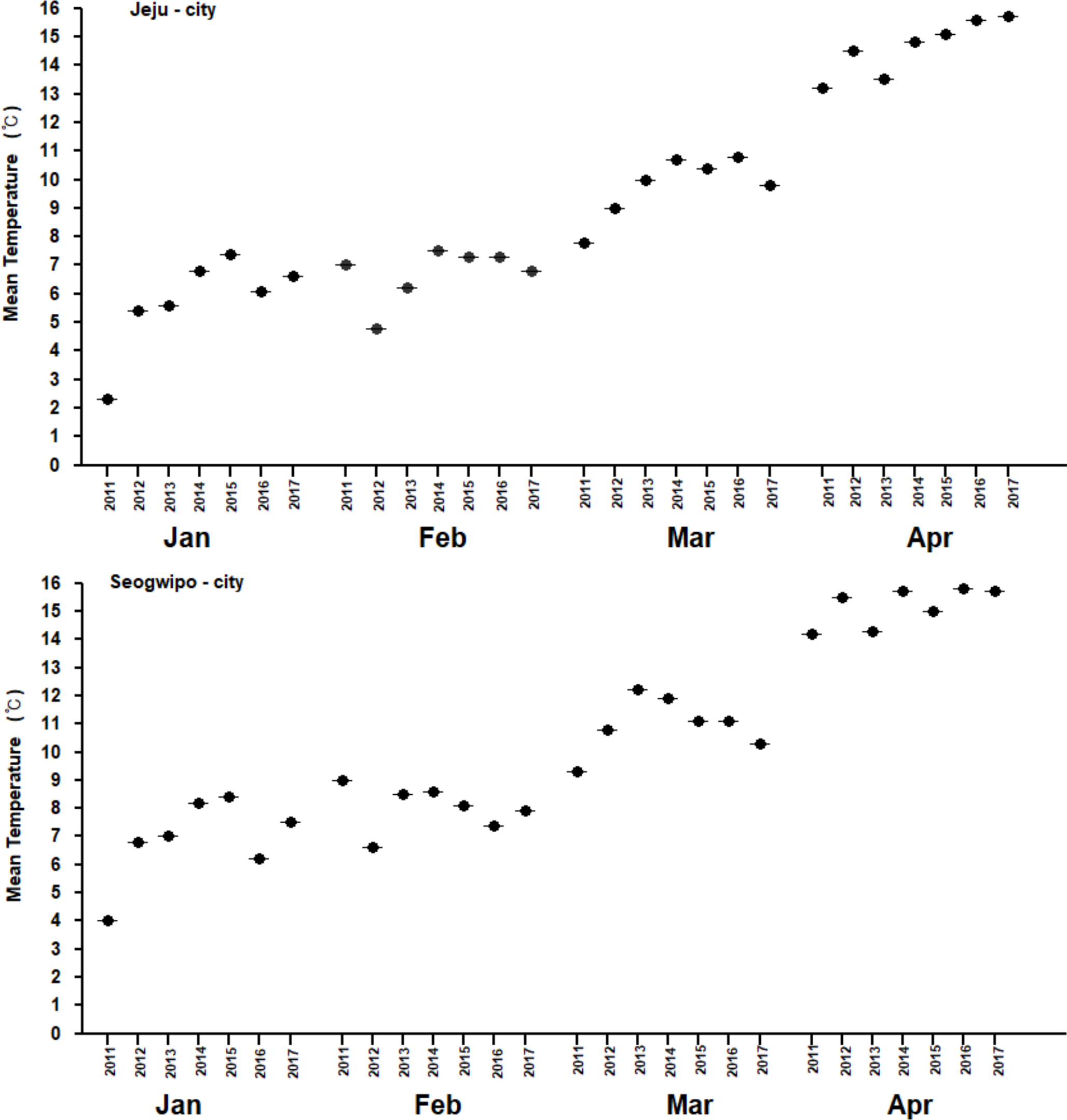
The monthly mean temperature during Japanese cedar efflorescence season in Jeju-City (NR) and Seogwipo-City (SR).

**Table 3.** The monthly mean temperatures (°C) during the JCP season (from January to April) in Jeju Island.

| Area | Year |  |  |  |  |  |  |  |  |
| --- | --- | --- | --- | --- | --- | --- | --- | --- | --- |
|  | Month | 2010 | 2011 | 2012 | 2013 | 2014 | 2015 | 2016 | 2017 |
| Jeju City (NR) | Jan | 5.3 | 2.3 | 5.4 | 5.6 | 6.8 | 7.4 | 6.1 | 6.6 |
|  | Feb | 7.3 | 7 | 4.8 | 6.2 | 7.5 | 7.3 | 7.3 | 6.8 |
|  | Mar | 9.3 | 7.8 | 9 | 10 | 10.7 | 10.4 | 10.8 | 9.8 |
|  | Apr | 11.8 | 13.2 | 14.5 | 13.5 | 14.8 | 15.1 | 15.6 | 15.7 |
| Seogwipo City (SR) | Jan | 7.5 | 4 | 6.8 | 7 | 8.2 | 8.4 | 6.2 | 7.5 |
|  | Feb | 9.3 | 9 | 6.6 | 8.5 | 8.6 | 8.1 | 7.4 | 7.9 |
|  | Mar | 10.8 | 9.3 | 10.8 | 12.2 | 11.9 | 11.1 | 11.1 | 10.3 |
|  | Apr | 13.3 | 14.2 | 15.5 | 14.3 | 15.7 | 15 | 15.8 | 15.7 |

**Table 4-1.** JCP sensitization rates among participants.

|  |  | 2010 year |  |  | 2016 year |  |  |
| --- | --- | --- | --- | --- | --- | --- | --- |
|  |  | Residential area |  |  | Residential area |  |  |
|  |  | Jeju City (NR) | Seogwipo City (SR) |  | Jeju City | Seogwipo City |  |
|  |  | Total (%) | Total (%) | Total (%) | Total (%) | Total (%) | Total (%) |
| JCP sensitization rates | Negative rate | 815 (89.8%) | 490 (76.4%) | 1305 (84.2%) | 479 (86.2%) | 385 (72.5%) | 864 (79.5%) |
|  | Positive rate | 93 (10.2%) | 151 (23.6%) | 244 (15.8%) | 77 (13.8%) | 146 (27.5%) | 223 (20.5%) |

**Table 4-2.** JCP sensitization rates in elementary school.

| 9 to 11 |  | 2010 year |  |  | 2016 year |  |  |
| --- | --- | --- | --- | --- | --- | --- | --- |
|  |  | Residential area |  |  | Residential area |  |  |
|  |  | Jeju City (NR) | Seogwipo City (SR) |  | Jeju City (NR) | Seogwipo City (SR) |  |
|  |  | Total (%) | Total (%) | Total (%) | Total (%) | Total (%) | Total (%) |
| JCP sensitization rates | Negative rate | 384 (92.80) | 164 (80.80) | 548 (88.80) | 251 (90.60) | 158 (79.80) | 409 (86.10) |
|  | Positive rate | 30 (7.20%) | 39 (19.20) | 69 (11.20) | 26 (9.40) | 40 (20.20) | 66 (13.90) |

**Table 4-3.** JCP sensitization rates in middle school.

| 13 to 14 |  | 2010 year |  |  | 2016 year |  |  |
| --- | --- | --- | --- | --- | --- | --- | --- |
|  |  | Residential area |  |  | Residential area |  |  |
|  |  | Jeju City (NR) | Seogwipo City (SR) |  | Jeju City (NR) | Seogwipo City (SR) |  |
|  |  | Total (%) | Total (%) | Total (%) | Total (%) | Total (%) | Total (%) |
| JCP sensitization rates | Negative rate | 279 (87.7%) | 174 (77.0%) | 453 (83.3%) | 130 (86.7%) | 129 (73.70%) | 259 (79.7%) |
|  | Positive rate | 39 (12.3%) | 52 (23.3%) | 91 (16.7%) | 20 (13.3%) | 46 (26.3) | 66 (20.3%) |

**Table 4-4.**
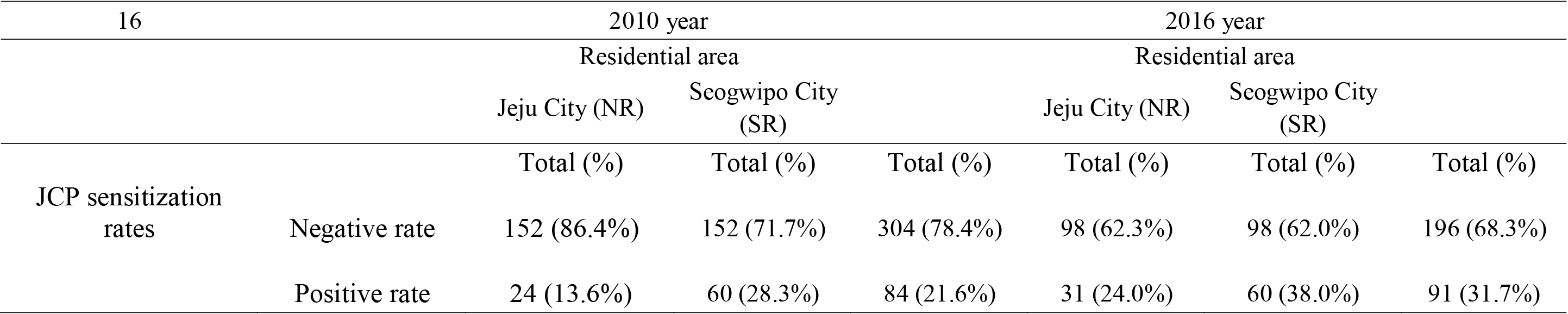
JCP sensitization rates in high school.

| 16 |  | 2010 year |  |  | 2016 year |  |  |
| --- | --- | --- | --- | --- | --- | --- | --- |
|  |  | Residential area |  |  | Residential area |  |  |
|  |  | Jeju City (NR) | Seogwipo City (SR) |  | Jeju City (NR) | Seogwipo City (SR) |  |
|  |  | Total (%) | Total (%) | Total (%) | Total (%) | Total (%) | Total (%) |
| JCP sensitization rates | Negative rate | 152 (86.4%) | 152 (71.7%) | 304 (78.4%) | 98 (62.3%) | 98 (62.0%) | 196 (68.3%) |
|  | Positive rate | 24 (13.6%) | 60 (28.3%) | 84 (21.6%) | 31 (24.0%) | 60 (38.0%) | 91 (31.7%) |

Our previous study showed that the JCP sensitization rate has been increasing each year, and the results of the present study further confirmed this trend (Table 4) [2].

Jeju Island is located at the southern end of the Korean Peninsula; this island is affected by climate change more than other regions of South Korea. The mean temperature of Jeju Island increased by 1.7°C from 1970 to 2011; during this period, the mean temperature of NR increased by 1.4°C, and the mean temperature of SR increased by 2°C [5].

JCP is the major aeroallergen contributing to the JCP sensitization rate and is the most prevalent SAR during the spring season in Jeju Island because the JC tree was systematically planted as a windbreak to decrease fruit loss and because JC is the dominant tree species.^2^ Climate change, such as increasing temperatures, causes earlier onset of the spring pollen season and of pollinosis caused by pollen [3].

## Discussion

In this study, we investigated JCP counts from 2011 to 2017, the monthly mean temperature during the JCP season and JCP sensitization among participants who were recruited among schoolchildren residing in 2010 and 2016 to understand the effects of increasing temperatures on the JCP count and sensitization rate.

The JCP season began earlier and lasted longer each year, the mean temperatures in January and April highly influenced the start and end of JCP production and the JCP season, and the JCP sensitization rate was higher in 2016 than in 2010. Therefore, our results show that increasing temperatures in January and April caused by climate change led to earlier and longer JCP seasons and that earlier and longer JCP seasons led to increases in the JCP sensitization rate and in SAR occurrence. We also suggest that there may be a significant increase in exposure to JCP caused by climate change and that further studies need to develop strategies to mitigate this exposure.

## Acknowledgments

This work was funded by a research grant from the Environmental Health Center (atopic dermatitis & allergic rhinitis) at Jeju National University, Jeju, Korea and the authors declare that they have no competing interests.

## Author Contributors

**Conceptualization:** Keun Hwa Lee.

**Data curation:** Keun Hwa Lee, Jung-Kook Song, Ju wan Kang.

**Formal analysis:** Keun Hwa Lee, Jung-Kook Song, Ju wan Kang, Sung-Chul Hong, Jeong Hong Kim, Dahee Park, Hye-Sook Lee, Jinho Jeong, Kyu Bum Seo.

**Funding acquisition:** Keun Hwa Lee.

**Investigation:** Keun Hwa Lee, Jung-Kook Song, Ju wan Kang.

**Methodology:** Keun Hwa Lee, Jung-Kook Song, Ju wan Kang, Sung-Chul Hong, Jeong Hong Kim, Dahee Park, Hye-Sook Lee, Jinho Jeong, Kyu Bum Seo.

**Resources:** Keun Hwa Lee.

**Supervision:** Keun Hwa Lee.

**Validation:** Keun Hwa Lee, Jung-Kook Song, Ju wan Kang.

**Writing - original draft:** Keun Hwa Lee.

**Writing - review & editing:** Keun Hwa Lee, Jung-Kook Song, Ju wan Kang.

